# Control mechanisms for stochastic biochemical systems via computation of reachable sets

**DOI:** 10.1101/079723

**Authors:** Eszter Lakatos, Michael P.H. Stumpf

## Abstract

Controlling the behaviour of cells by rationally guiding molecular processes is an overarching aim of much of synthetic biology. Molecular processes, however, are notoriously noisy and frequently non-linear. We present an approach to studying the impact of control measures on motifs of molecular interactions, that addresses the problems faced in biological systems: stochasticity, parameter uncertainty, and non-linearity. We show that our reachability analysis formalism can describe the potential behaviour of biological (naturally evolved as well as engineered) systems, and provides a set of bounds on their dynamics at the level of population statistics: for example, we can obtain the possible ranges of means and variances of mRNA and protein expression levels, even in the presence of uncertainty about model parameters.

## Introduction

Much of the research in synthetic and systems biology in the last decade has focused on the study of elementary biological systems. This has often been with the aim of controlling and modifying them in order to achieve new functional modules exhibiting novel and useful behaviour [1], such as sustained oscillations [2] and bistability [3]. It is now becoming possible to use engineered biological systems made of characterised components to solve specific problems, such as information processing, energy production or production of chemicals.

However, biological systems are inherently noisy and probabilistic in nature, which can pose significant difficulty for one aiming for a reliable, well-characterised module. Although several external control techniques have been developed which are able to avoid some of the variability in a population [4, 5], noise at the level of molecular processes is often unavoidable and does, for example affect quite profoundly how information is transmitted along the molecular networks underlying cell function [6, 7]. Furthermore, due to the unreliability of measured quantities, our understanding of the underlying mechanisms might be mistaken leading to sub-optimal analysis and design. Therefore we have to assess the practical limits on the amount of noise in a general biological module, in order to evaluate the efficiency of a control design or the reliability of a mechanistic model.

Reachability analysis has been widely used in control design and engineering for applications such as verification of electrical or mechanical networks and hybrid automata [8, 9]. The analysis focuses on the computation of the subset of the state space that can be reached within a certain time-limit, given some starting position of the system and external inputs. The technique can also be used to verify that a certain undesired state is not reached under realistic operating condition, or that the behaviour of the system is robust and qualitatively/quantitatively holds for different conditions, including different realistic inputs to the system.

To this end reachability is generally calculated under varying levels of uncertainty regarding details of the system – such as initial state, input signal and rate parameters — usually formalised by assuming that these parameters come from a set of plausible values. Therefore, unless analytical solutions are derived for some abstraction of the system, the applicability of the computation heavily relies on the choice of the set representation. Some representations might prove computationally expensive and hence impractical for high dimensional systems, while a simple shape representation can lead to crude over-approximation of the reachable set. Methods have been proposed using several techniques such as polygonal projections [10], oriented hyper-rectangles, special polyhedra [11], ellipsoids [12], or level sets [13]. In this work we use zonotopes [14], a centrally symmetric type of polytopes that can be conveniently represented by a list of vectors.

Although there is already an enormous body of work on reachable set computation for problems in engineering, including highly nonlinear cases [15], hybrid automata [16], and differential-algebraic equations [17], the complexity, frequent nonlinear behaviour, and strict constraints on many of the model parameters of biochemical systems require the development of specialised analysis methods. There are already a few applications of reachability techniques to biological examples with special emphasis on the treatment of the non-linearity of the system, either through direct computation [18] or through hybridisation-based methods using either static or dynamic partitioning of the state space [19, 20, 21]. The work by Dang et al. [21] has also been expanded to take into account the possible lack of knowledge of parameter values [22]. The stochasticity in biological systems has been tackled in even more diverse ways: through computing bounds on the probability function [23], using stochastic hybrid systems [24], or analytically deriving invariant sets for linear equations obtained from the stochastic model [25].

In this work we propose a computationally efficient and flexible method to compute the reachable set (in a zonotopic representation) of stochastic biochemical systems; besides stochasticity we also consider possibly nonlinear rate laws, controlled or uncertain input signals, and uncertainty about model parameters that might also represent control over these values. The main steps behind our derivation are the following: (i) we first obtain an ODE representation of the system’s mean and (co-)variances; (ii) then use an iterative procedure to obtain a conservative approximation of consecutive reachable sets up to a final time of interest. Here we primarily use the Linear Noise Approximation (LNA) for the first step, but also present the Moment Expansion Approximation [26, 27, 28] to demonstrate that other methods with different applicability can be equally used to generate equations for the second step. We derive a new method to tightly approximate realistic biological input signals, and a piece-wise temporal linearisation method is applied to deal with common nonlinearities. We also give conservative approximation formulae for the reachable set if rate parameters of the system are not precisely known — which is very often the case in biochemical systems. The method is demonstrated on two elementary modules fundamental to mathematical models and regulatory designs of biochemical networks. The first system presents the use of reachability analysis for the study of noisy biochemical reactions and evaluating a control on the levels of cell heterogeneity; the second example considers the task of model (in)validation for cases when high cell-to-cell variability poses a challenge to estimating the system’s true behaviour.

## Methods

### Zonotopes

A zonotope [29] is described by the position of its centre (*c*) and a set of generator vectors (*g*_1_,…,*g*_*p*_)as

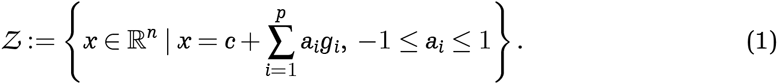

In the followings we use the shortened notation (*c*; *G*) to represent 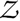, where 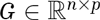 is a generator matrix formed from the generator column vectors. Zonotopes are a convenient representation as they are closed under Minkowski-addition and affine transformations, the two key operations in reachability analysis [14]. Furthermore, the above can be calculated through simple matrix-vector operations: given zonotopes 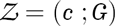 and 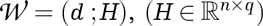 and the affine transformation 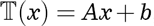,

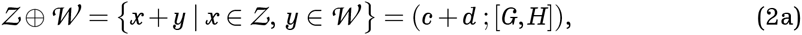

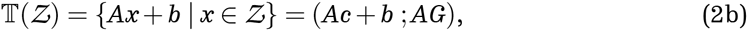

where [*G,H*] is the concatenation of the ‘vector lists’.

### Linear Noise Approximation

Given a stochastic system with *N*-dimensional state variable 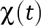 describing the abundance of modelled species at time *t*. We divide the state variable into a macroscopic part, and random fluctuations, as 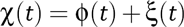. If the system is defined by a stoichiometry matrix (*S*) and a collection of reaction propensities, 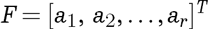, the corresponding equations of time-evolution are,

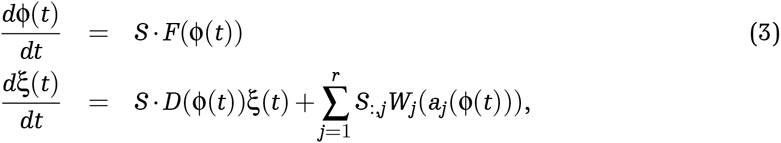

where *W*_*j*_(*d*) is a Wiener process and 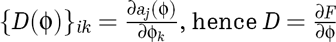 is the Jacobian of the system. In applications where population-level behaviour is of interest Eq. (3) can be used to obtain equations that describe how the mean and variance of the probability distribution of 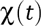 changes. The decomposition of the state variable makes it straightforward to follow the change in the mean of the system, as it is already given by Eq. (3); while the evolution of the covariance matrix (Σ) is calculated as

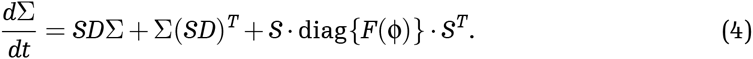

Thus a set of ordinary differential equations can be derived that follows the time-dependent change of the mean values and (co)variances of all species, summarised in the 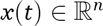 vector, where *n* = *N*(*N*+3)/2:

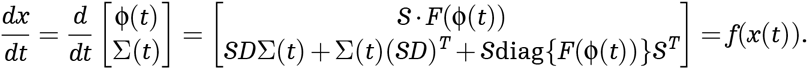

### Moment Expansion Approximation

The Linear Noise Approximation is based on the assumption that the system noise is well described by a normal distribution, and molecules are present in at least moderately high amounts. In many cases it offers a good approximation even when these conditions are not met. However, to handle more problematic cases, we also use Moment Expansion Approximation, a less constrained moment generating method, to obtain ordinary differential equations. A user-friendly Python-based interpretation for automatic computation of this step is available from github [28].

In brief, given the aforementioned stochastic system formed by 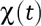, *S* and *F*, we obtain the moment generating function of the species’ probability distribution, 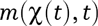, which is then differentiated with *k* times to lead to equations of the *kth* order moments. These expressions are evaluated with the help of two consecutive Taylor expansions. As for nonlinear systems the resulting equations would be in theory infinite for – due to moments always depending on subsequent moments – we finish by applying moment closure formulae (*mc*^*k*^) to substitute the highest order terms 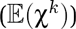 with expressions of means (ϕ(*t*)) and (co)variances (Σ(*t*))

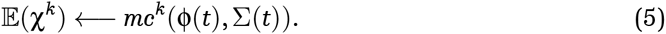

A detailed description of the algorithms and applicability of moment expansion can be found in [26] and of closure methods in [27].

For reachability analysis our primary interest is in the mean and variance of molecules of the system, and Moment Expansion can be used in three ways to derive these. We can (i) set the expansion to be only up to second-order moments; (ii) use the step in Eq. (5) to substitute all higher order moments, not just the highest order ones; or (iii) use the whole system of *k* moments as our state vector, to make sure no essential influence on means and variances is omitted.

### State-space representation

In the next step we consider a general deterministic system, *S*, described by the evolution of the *n*-dimensional state variable, *x*(*t*)

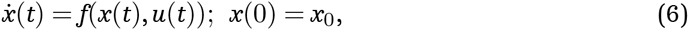

where 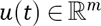 is an input signal, and *f*(*x*, *u*) is a generally nonlinear but time-independent transition function. We focus on the case where the input-dependence in the above equation can be separated as

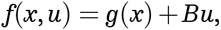

where 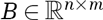 is the input matrix, which specifies which states are affected by the inputs. We further assume that instead of *x*_0_, a set containing all possible initial values in the state-space, *I,* is given – for example because we observe a range of expression values in a population of cells. Similarly, the input signal also comes from a set, *U,* which is bounded by some value 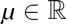 so that 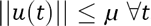. Then Eq. (6) is presented as a differential inclusion [30] of the form

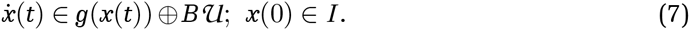

The reachable set of this system at any time *t* is defined as

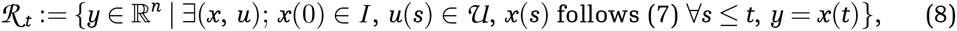

i.e. all the states we can achieve by having the above described system start from a possible initial state and affected by plausible input signals. Similarly, we can define 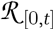 as all the states reachable within the time interval [0,*t*], by computing all individual reach sets: 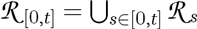.

### Reachable set computation

We start our derivation considering the case when function *f* is a linear function of *x;* under such condition the system can be represented in the linear time-invariant (LTI) form,

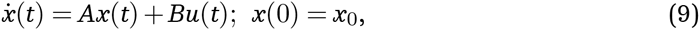

where 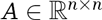 is the state (transition) matrix. We focus on single input systems, where *B* is an *n ×* 1 matrix; however, all results can be easily generalised to other values of input dimension, *m.* The solution of this system is generally given as

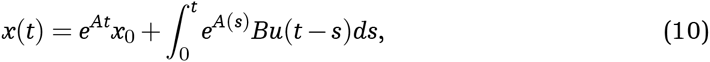

which can be further simplified through assuming a constant input signal and an invertible transition matrix

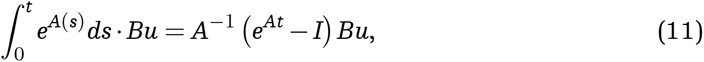

where *I* is the *n × n* identity matrix. In the followings for convenience we use the notation

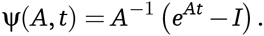

We consider an equidistant partitioning of the time horizon into *N* intervals, with time-step τ = *T/N.* The base of the algorithm is the propagation of reachable sets from *I* to 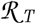 in the autonomous system, i.e. when *u* ≡ 0. Given a reachable set at an arbitrary time-point, 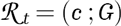, the new set reached under zero input can be calculated according to the first element in Eq. (10). This equals the affine transformation 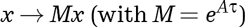 and hence, according to Eq. (2b), the new reachable set is given by the zonotope 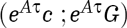. The same dynamics hold for all *t,* and the transition matrix can be iteratively applied to propagate *I* up to the final reach set, 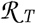. Furthermore, functions of the transition matrix, such as 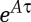, have to be calculated only once through the initialisation of the algorithm.

However, in systems with uncertainty (that might be of the form of an unknown input signal) we need to transform and enlarge, or *bloat,* the set 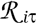 in each iteration, to obtain all states the system can take under an admissible input in 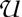. This is done by taking the Minkowski sum of 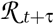 and bloating sets, β_*μ*_ and β_*δ*_, corresponding to uncertainty in the input and parameter values, respectively.

In case the aim is to map the entire space the system explores between times 0 and *T,* the reachable sets of the time intervals, 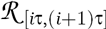 has to be derived from the two end-points. Generally, this can be done by computing the convex hull of 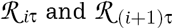, and then “bloating” this set to contain all affine solutions for one time-step (see [31] for an illustration). In order to avoid the exact computation of a convex hull, as zonotopes are not closed under this operation, a conservative (“over–”)approximation is presented in [14] — we use this approximation to derive a set enclosing all points reachable between 0 and *τ.* All following intervals can be propagated from this set, as for all 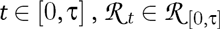 and hence 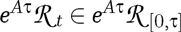. To summarise, using the short form 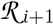 to denote either 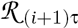, or 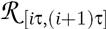, depending on the type of analysis we aim for, given 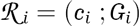, the next reachable set is calculated as

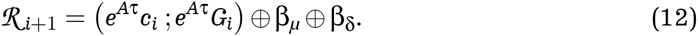

### Input for biological systems

In the algorithm proposed by Girard [14], the input set, 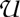 is taken to be an *n*-dimensional hypercube enclosing all points between [−*μ,μ*] in all dimensions: equivalent to the radius *μ* ball in the infinity norm. Such a generalised approximation of the input set is sometimes necessary, as the analysis is motivated by a target or an avoidable set in the state-space and the input set is meant to capture all variations induced by noise in a physical/electrical system.

In the case of biological inputs, however, we usually have more detailed information about 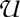, meaning that the aforementioned general input set would contain many implausible signals, and hence the over-approximation would be too loose to provide useful information on the actual reachable sets. For example, in the typical case the control input is implemented as a certain type of molecular species added to the system; this cannot take negative values, and it is also reasonable to assume an upper limit on the number of molecules injected at any time point, which can be viewed as the bound *μ.* Furthermore, the input matrix, *B,* determines how much each variable is effected by the input signal and hence cannot be neglected.

As 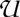 is typically not centred around 0, we divide the input effect into a drift term, corresponding to the effect of the centre of 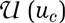, and an uncertainty term representing our lack of knowledge about the exact input value, i.e. the maximal difference, *u*_*d*_, between the centre and possible values of 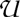. Thus the whole input set is taken into account as 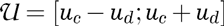; for instance, in the example above, *u*_*c*_ = *μ*/2 and *u*_*d*_ = *μ*/2. The drift and uncertainty terms are calculated using equation (11), and the bloating set will be the zonotope

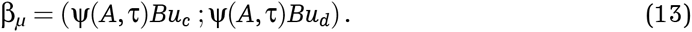

This calculation in this exact from is only possible if *A* is invertible — in the singular case we can use an approximation of Eq. (11), based on the integral of the Taylor-series of 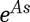 and compute a bloating factor 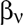 to correct for the small error thus introduced [17]:

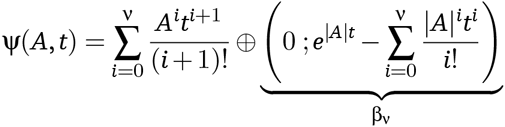

### Parameter uncertainty

Often we have to make predictions but only have approximate values of the parameters contributing to matrix *A*, either due to imperfect knowledge of reaction rates [32], or because we can control a reaction through some interaction that is not or cannot be modelled explicitly. Therefore, we derive a way to account for uncertainty in case the matrix *A* is also drawn from a set or ensemble of matrices. For example, we can consider the case where a single reaction rate, *k,* possibly effecting more than one element of *A*, is not known precisely or controlled. We assume that *k* has some plausible upper and lower bounds and hence it can be considered as coming from an interval centred at the nominal value 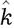, i.e. 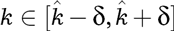. We approach the problem by following the nominal dynamics using the previously defined matrix, *A*, and defining a bloating term to enclose all solutions arising from admissible parameter values, so that

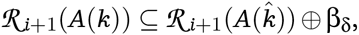

where *A*(*k*) denotes the matrix formed using the parameter value *k.* For a positive *x,* we can bound *A*(*k*)*x* with 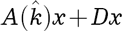 where *D* is the *n × n* matrix computed as 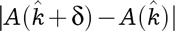. To make *Dx* independent of the state variable — and thus derive a general formula –, we use a conservative estimation of 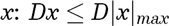, where |*x*|_*max*_ can be approximated generally for the whole algorithm, or, more practically, in each time step based on the states in the current reachable set. We derive |*x*|_*max*_ from the zonotope definition in Eq. (1): given a reachable set in the form (*c*, *G*), the *i*th coordinate of the maximal vector is computed by the formula

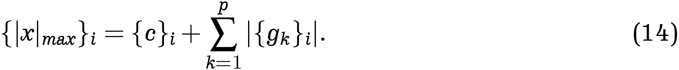

Note that |*x*|_*max*_ might not be in the reachable set, but there is at least one point for each coordinate for which {*x*}_*i*_ = {|*x*|_*max*_}_i_ holds. |*x*|_*max*_ is derived as |*c*| + ∑|*G*| with all summations carried out *by rows*. We also take into account a rough estimate of the next reachable set, so that in non-converging cases |*x*|_*max*_ is not underestimated: 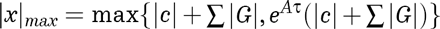.

For each coordinate of vector *D*|*x*|_*max*_ the difference between dynamics under any *k* and the centre, 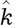, can be enclosed in the zero-centred set:

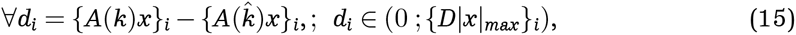

from which a zonotope considering all coordinates can be obtained as (0; diag{*D*|*x*|_*max*_}). From here we proceed considering this zonotope as an input set centred at 0 – in agreement with the fact that trajectories of the nominal value, 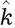, are already calculated through the transition matrix. Therefore the bloating set accounting for parameter uncertainty, (β_δ_, can be computed as

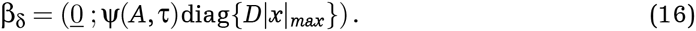

### Nonlinear systems

Biological systems are often nonlinear, as even the simplest dimer-formation requires second-order rate laws that cannot be eliminated from the system. Nonlinear systems can lead to complex and unpredictable behaviour and their analysis can be difficult. The most popular way to overcome this is by performing a (piece-wise) linearisation of the system, in each segment of an either temporal or spacial partitioning of the time-horizon and state-space studied. In our work we apply the former approach, and at each step linearise the system around its current centre using first-order Taylor expansion, as described in [33]. This results in a system which is piece-wise linear, and has LTI properties between two time-steps, but overall time-varying and is only an approximation of the real underlying dynamics. The linearised system calculated at time *i*τ is

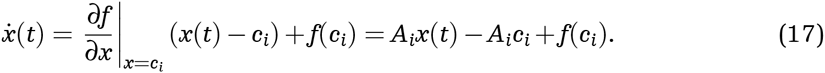

This system can be used to obtain the next reachable set together with a bloating factor that accounts for the difference between the original and the linearised dynamics

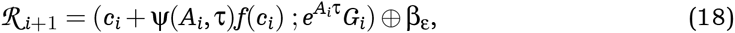

where β_ε_ can be computed iteratively as proposed in [33]. We use the zonotopic set representation to obtain a tight estimate of the effect of nonlinearity Consider the error function ε(*x*) = |*f*(*x*) – [*A*_*i*_(*x* – *c*_*i*_) + *f*(*c*_*i*_)]| – using a substitution of *x* by (*c*_*i*_ + *dx*), we obtain ε as a function of *dx,* the distance between a point and the centre. We define an upper bound for *dx,* as in Eq. (14): 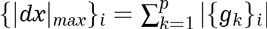, and approximate ε(*x*) with a function monotonously increasing with *dx.* Hence we obtain an estimation for which 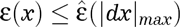 and construct a generator set

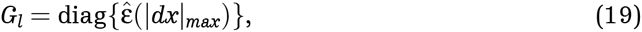

which represents a bound on the nonlinearity of the system.

To obtain the reach set of the time-interval [*i*τ, (*i* + 1)τ], further approximations have to be applied to enclose all trajectories between the two time-points. As the piece-wise linear system is time-varying, the estimation of the time-interval [0, τ] is not sufficient. Instead, as mentioned before, the convex hull of two delimiting reach sets is calculated and bloated, the bloating factor for which can be derived in several ways, e.g. as in [14] or [33]. Alternatively, we could replace the linearisation with a more flexible description of the stochasticity than the LNA, such as moment closures [27] or finite–state projection methods [34].

## Example Applications

### Control of gene expression noise

The first example considered is controlled stochastic gene expression system [35, 25]. Despite its simplicity, this model is one of the most important examples in practice as protein production is a necessary and elementary building module in real and synthetic biological systems. Many applications might rely on the steady operation of such a unit, hence it is of great practical interest to know what the achievable noise levels and possible states of operation are. The original stochastic model has two variables (mRNA and protein copy numbers, *m* and *p* respectively) and four reactions (transcription, mRNA degradation, translation and protein degradation), characterised by the set of reaction propensities and stoichiometry matrix

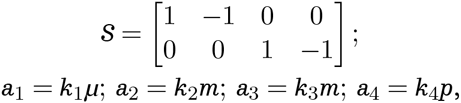

where the multiplier in the first reaction, *μ* represents our control over the transcription rate, assuming either discreet (*μ* = 0 or *μ* = 1) or continuous (*μ* ∈ [0,1]) values. A typical control signal would be a sequence of zeros and ones, switching transcription on and off for some time period. After performing the LNA, we obtain a set of five ordinary differential equations, determining the time evolution of the mean of *m* and *p*, the variance of *m,* the covariance of the two species and finally the variance of the protein abundance. The system is started from 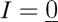 and we investigate the reachable sets up to time *T* = 10 minutes with time-step τ = 0.01 and reaction rates 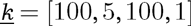.

Figure 1 shows projections of the final reachable set together with two sample trajectories. The samples are computed as the mean and variance of the population consisted of 10000 direct realisations of the original stochastic system. Input signal *μ*(*t*) is defined as a piece-wise constant function with values randomly drawn from {0,1} and switching every 60 or 30 seconds (*u*_1_ and *u*_2_, respectively); the signal is kept the same for all realisations contributing to a particular trajectory in fig 1. We also take into account some uncertainty regarding the protein degradation rate, i.e. that the value of *k*_4_ is unknown or controllable to within 5% of the nominal value, 1. The light blue regions in figure 1 show the estimate of the reachable set under such uncertainty. Interestingly, while a substantial part of the mRNA-protein mean space can be covered, the reachable states on the protein mean-variance plane are limited to a very narrow band. Therefore, if relying on the production of a protein with this module we will have to make a compromise: either have a low number of proteins produced, or a high amount but with great variability (as may be expected given the Poisson nature of this process).

**Figure 1:**
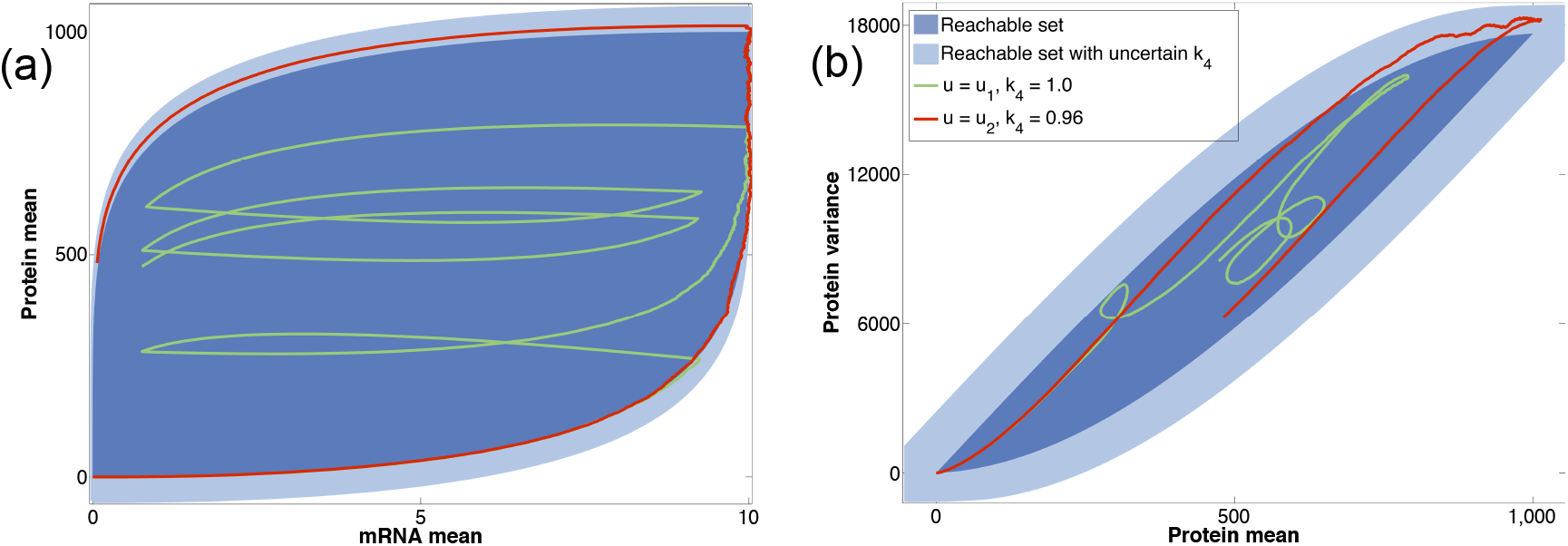
Reachable states of the stochastic gene expression system with controlled transcription and additional uncertainty. Blue shaded regions show projection of the final reachable set to (a) the mRNA mean-protein mean plane and (b) the protein mean-protein variance plane. Dark and light blue shades indicate reachable sets without and with 5% uncertainty in parameter *k*_4_. Red and green example trajectories are calculated from 10000 exact simulations, under the input sequences *u*_1_ = [1,0,1,1,1,0,1,0,1,0] and *u*_2_ = [1,1,1,1,1,1,1,1,0,0,0,0,0,1,0,1,1,0,1,1], respectively, with protein degradation values as indicated in legend.

### Model validation

In the our second application we consider a chain of three molecules effecting each others production and aim to demonstrate how our reachability analysis can contribute to model validation [36] for stochastic systems. The molecules A, B and C in Figure 2 can represent any chain of interacting species with similar reaction networks; for example a simple model of transcription factors, where only protein levels are modelled explicitly. The regulatory effect of these molecules is through the production rate of their target species. Three different wiring schemes are considered: (i) molecule A induces molecule B that in turn activates the production of C; (ii) molecule A also activates molecule C, such that this effect is more profound than the activation via B; (iii) molecule C feeds back and induces the production of A. In each model we simplify the mathematical description by assuming equal degradation rates for all species; hence the models can be summarised by 5 parameters: 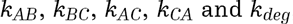, where the subscript *XY* refers to the activation of *Y* by *X*. In all models 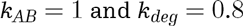 and the other parameters are chosen to reflect the connections of the model and produce similar maximal values in the output, C.

**Figure 2:**
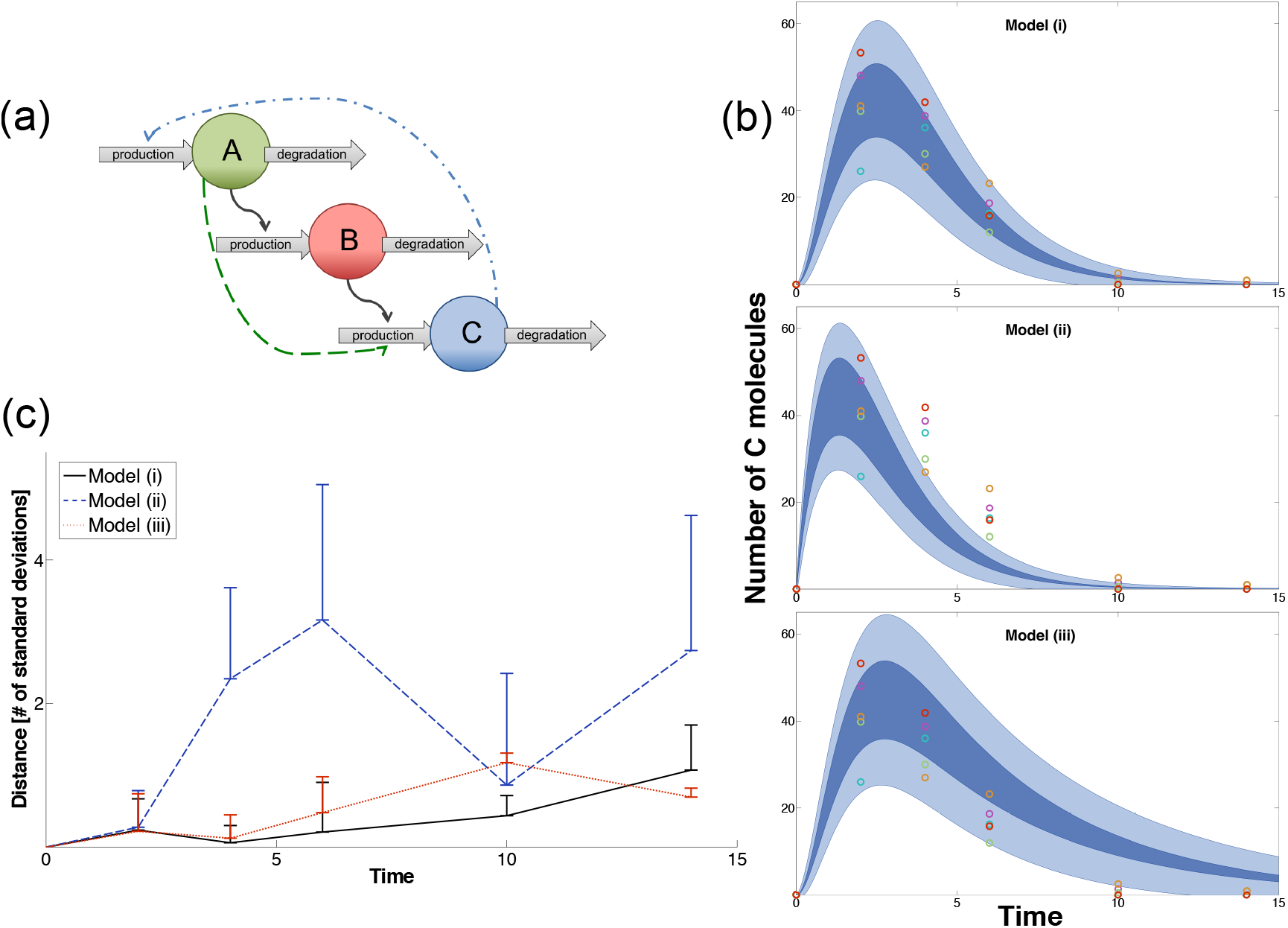
Model evaluation by comparing reachable sets and single measurements. (a) Schematics of the three reaction chain models. Models differ in rates corresponding to dashed arrows. (i): *k*_*BC*_ = 1, *k*_*AC*_ = *k*_*CA*_ = 0. (ii): *k*_*BC*_ = 0.1, *k*_*AC*_ = 0.9, *k*_*CA*_ = 0. (iii): *k*_*BC*_ = 1, *k*_*AC*_ = 0, *k*_*CA*_ = 0.2. (b) Reachable region over time of the output (molecule C) starting from the initial set 80 ≤ *A*_0_ ≤ 120, *B*_0_ = *C*_0_ = 0. Dark areas represent reachable values of the mean, light blue shades are the *±* 1 standard deviation region computed with the maximal reachable value of the variance. Coloured circles are sample points taken from single exact simulations of Model (i). (c) Distance of observation points from the reachable set of mean values. Lines show the average distance of observed data at each time point for each model (colours as indicated in legend), the top of error bars depict the maximal distance at the evaluation points.

In order to model measurements, we generate five individual stochastic trajectories from Model (i), with parameter values 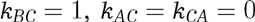 and an initial A value randomly chosen from the range [80, 120]. We then sample the amount of molecule C – as the assumed output is experimentally measurable – from these simulations at six time-points; and evaluate the distance of each measurement point from the reachable set of mean values at that specific time. The distance is normalised by the maximum reachable value of standard deviation and zero for all points within the reachable set. As Figure 2(b)-(c) shows, it is very unlikely the observations would arise from Model (ii). Although the true model shows the best correspondence to the data, Model (iii) — representing a feedback system — cannot be discarded due to the wide range of values molecule C can reach in that specific wiring scheme. If the relative noise level is reduced by raising the average initial abundance (to ~ 300 molecules), or measurements from another species (e.g. molecule A) is also available, the distinction between different models becomes clear with Model (ii) unambiguously fitting the data best.

### Discussion

In this work we have introduced a method to compute the states a stochastic biochemical network can access under a control input signal. The method is particularly applicable to determine reachable mean-variance values of the investigated species, and hence estimate noise levels of a system. Our aim is to provide a computationally efficient tool that could be used in the search for the best modelling description or regulatory design of stochastic networks.

Here we used the Linear Noise and Moment Expansion Approximations to extract deterministic equations of the mean and covariance matrix that served as a basis for our reachable set computation. Both can be accessed on github as parts of our Python package, MEANS [28]. Then we have presented a general algorithm for linear and nonlinear reachability analysis based on zonotopes, together with formulae for handling relevant input types: addition of molecules and control of a reaction rate. All steps are implemented in Matlab to semi-automatically generate the reachable sets for a problem defined in terms of a transition function, input matrix and bounds, and parameter uncertainty. The algorithm can be executed on any given set of ODEs describing some characteristics of the system, hence it is applicable regardless of the approximation method used for the generation of equations and also to deterministic biological models.

We demonstrate the method on two schematic models of biological macromolecules: a controlled gene expression system and a cascade of modifying molecules, e.g. transcription factors. In both examples the approximation is conservative, hence trajectories randomly generated from a set of admissible signals and initial values are confined within the set our method predicts. In linear cases, if we have precise knowledge of the parameter values and input bounds, the predicted reachable set will be exact as well, i.e. each point in it can be actually accessed by an appropriate input sequence. This cannot be ensured for nonlinear systems (or uncertain rate values) as the conservative estimates used for the nonlinearity (or reaction rate effect) are obtained from approximations the system might note take. However, our estimation reflects the general characteristics of the system – e.g. if some variables of a generally nonlinear system have linear equations – and hence unnecessary over-approximations are avoided to provide a tight estimate. This is crucial in the evaluation of various control designs on the basis of mean-variance values reachable through them.

Our method is based on a zonotope representation of reachable, initial and parameter sets. Set operations can be efficiently computed using this representation method, which allows us to apply our reachability analysis to many-variable, complicated biological networks. However, the number of generator vectors in the reachable set increases in every iteration, and for a long time horizon or small time step the algorithm can become expensive regarding memory space. In smaller systems, such as our examples, this effect is still negligible, but for high-dimensional cases the increase is more significant. For such cases, one can turn to the zonotope-reduction technique presented by Girard [14] to limit the size of zonotopes to an adjustable value.

Note also that zonotopes, and hence the sets in our analysis, are by definition convex and centrally symmetric. Therefore systems where the actual reachable set is concave or consisted of multiple sets — especially networks with bi- and multi-stability — will be largely over-approximated. We tested the algorithm on a bistable model of stem cell differentiation. As the two expression level states in such cases are typically of different magnitudes, our method is forced to enclose a high proportion of negative values besides also incorporating a non-accessible region between the two states. However, we also found that this drop in approximation quality is a good indicator of the system entering a bistable regime of initial conditions/input signal. A similar over-approximation case can arise for imprecise parameter values of the transition matrix: although admissible parameter sets are symmetric sets given as a nominal value with error bounds, the influence of parameters is usually not symmetric on the reachable set. Therefore zonotopes lead to a rough approximation when some rates show high uncertainty. Just like in the case of multi-stability, a preliminary analysis using our general method can reveal these issues, and point one towards a more exhaustive study using a series of reachable sets for fixed values (or small intervals) of initial values or rate parameters to cover the range of interest and extrapolate for tighter approximation.

As often the case in the analysis of biochemical systems, incomplete information about the model structure and crucial constants of the system makes a detailed and informative analysis impossible. Our method is not applicable for the exploration of differences in model structures; this problem can be treated implicitly by either doing independent analysis of all candidate models or choosing rate parameter uncertainty such that zero (i.e. no interaction) is amongst the admissible values. The latter is likely to give rise to an over-generalised approximation, as described above; while the first one is only advisable for a small number of possible models, like in our second example. Furthermore, limited measurements and high levels of noise can also influence the method’s power for validation, as Figure 2(c) demonstrates: in such cases flexible models might be favoured even over the true model. Therefore we advise the use of other tools, such as Topological Sensitivity Analysis [37] coupled with our method for a more thorough investigation of model space.

## Acknowledgment

EL acknowledges support from the Schrödinger Scholarship; MPHS is supported by the BBSRC and through HFSP grant RGP0061/2011.

